# Regulator-derived growth and fitness costs of *Salmonella* SPI-2 expression are environment specific and T3SS independent

**DOI:** 10.64898/2026.04.10.717833

**Authors:** Madison Spratt, Cece Stumpf, Mia Vigil, Karen Snyder, Keara Lane

## Abstract

Bacterial pathogens must evolve regulatory mechanisms that balance the necessity of virulence genes with the costs associated with their expression. *Salmonella* Pathogenicity Island-2 (SPI-2) is a genomic locus that encodes a Type 3 Secretion System (T3SS) and secreted effectors necessary for intracellular survival and replication. Recent work has shown that SPI-2 is expressed heterogeneously, suggesting uniform or aberrant expression of the system carries fitness costs. Here, we report a negative correlation between microcolony growth and SPI-2 expression, suggesting a growth cost to expression of SPI-2. To quantify and mechanistically evaluate this cost, we employ knockout and synthetic expression of the SPI-2 master regulator *ssrB* to eliminate or force the expression of SPI-2 genes. We find that *ssrB* expression causes deficits throughout the growth curve and puts cells at a competitive disadvantage in acidic, nutrient limited media. We further show that these deficits are environment specific, and not observed until stationary phase in neutral, rich media. We also observe unexpected cell morphology effects of *ssrB* expression, suggesting its effects on the cell extend beyond its regulation of SPI-2 genes. Using known *ssrB* mutations we find that DNA-binding activity, but not phosphorylation, of SsrB is necessary for growth costs. Lastly, we show that this growth cost is incurred even in the absence of the SPI-2 genetic locus, indicating that it is inherent to expression of *ssrB*, rather than its primary downstream virulence gene targets. These results demonstrate that SPI-2 expression is coupled to context specific cell growth and fitness deficits induced by its master regulator, offering a potential benefit to heterogeneous expression.

**SIGNIFICANCE:** Virulence factors are often heterogeneously expressed in bacteria, offsetting the resource and energetic burdens they impose. Characterizing such burdens reveals the evolutionary pressures that act on virulence genes and uncovers the logic behind regulatory mechanisms that drive heterogeneity in their expression. This study evaluates the growth and fitness costs associated with expression of *Salmonella* Pathogenicity Island-2 (SPI-2), which encodes a Type 3 Secretion System and secreted effectors required for intracellular survival and replication. The expression of the SPI-2 master regulator *ssrB* is shown to impose environment-specific growth and fitness deficits. These deficits persist in the absence of T3SS genes, suggesting that costs originate from SsrB acting on other target genes. These results provide critical context to recent observations of heterogeneous SPI-2 expression, elucidating the fitness cost this heterogeneity has evolved to offset.

## INTRODUCTION

The ability of a bacterial pathogen to colonize a host and cause disease is dependent on virulence factors. By definition these factors are essential for a pathogen’s lifecycle; however, this necessity must be balanced at the regulatory level with the energetic and resource burdens that they impose on a bacterium (Davis, 2020). Directly quantifying the costs associated with aberrant expression of a given virulence system and evaluating how these costs are incurred reveals the selective pressures that have acted on it, providing insight into the logic behind its regulatory mechanisms.

A common strategy employed by pathogens to manage the cost of virulence factor expression is phenotypic heterogeneity, in which subpopulations of expressors and non-expressors arise from the same clonal population (Davis & Isberg, 2019; Schröter & Dersch, 2019). Heterogeneous expression of virulence factors has been widely observed, including in Type 3 and 6 secretion systems, flagella, and toxin and biofilm matrix production (Hautefort et al., 2003; Kerwien et al., 2025; Lin et al., 2021; Koirala et al., 2014; García-Betancur et al., 2017). The benefit of heterogeneous virulence expression is generally assumed to be that energetically costly macromolecular production and function are limited to a subset of the population. This heterogeneity can confer a bet-hedging-like benefit, in which variable expression maximizes population survival in the presence of ambiguous signals (Ciolli Mattioli, 2026), or a division-of-labor benefit, in which functional differentiation is necessary for population-level infection success (West & Cooper, 2016). While experimentally supporting either of these mechanisms from an evolutionary standpoint is challenging in most systems (de Jong et al., 2011), quantifying the cellular burden of virulence factor expression allows for rational inferences about the evolutionary advantages of phenotypic heterogeneity.

While the cost of virulence factor expression is often attributed to the production and function of resource-consuming protein complexes, this generalization is complicated by several factors. First, for such a cost to be evolutionarily significant, cellular resources must be sufficiently limited in the pathogen’s native infection environment such that virulence expression causes a measurable growth or fitness deficit. Given the diversity of stress conditions, nutrient availability, and environmental stimuli a pathogen experiences over the course of an infection, growth and fitness costs of virulence expression may only be observed in specific contexts, or perhaps not at all, as has been shown in Type 6 Secretion System (T6SS) expression in *Vibrio fischeri* and *Escherichia coli* (*E. coli*) respectively (Septer et al., 2023; Taillefer et al., 2023). Second, while virulence structures, such as secretion systems, may be conserved across species, the regulatory mechanisms activating them have evolved in dramatically different environments. This can make the fitness cost of virulence factor expression challenging to disentangle from an environmentally co-regulated gene expression program. Examples of this include the coregulation of the stringent response with activation of *Legionella* invasion genes during its transmissive lifecycle phase (Oliva et al., 2018) and the growth rate-dependent regulation of flagella in *E. coli* (Sim et al., 2017). Lastly, the fitness costs of virulence factor expression are not limited to growth deficits. Host innate immune receptors broadly detect flagellin and T3SS components, imposing an additional selective pressure against their expression (Miao et al., 2006; Molofsky et al., 2006; Franchi et al., 2006; Miao et al., 2010). Given this complexity, it is critical to assess costs of virulence factor expression in a system-specific, context-dependent manner.

We have recently characterized bimodality in the expression of *Salmonella* Pathogenicity Island-2 (SPI-2) (Spratt & Lane, 2025), a virulence factor encoding a T3SS and secreted effectors required for the intracellular survival and replication of *Salmonella enterica* serovar Typhimurium (STm). Upon entry into host cells, STm activates SPI-2 to translocate effector proteins into the host cell cytosol, which repress innate immune function and promote the stability and maturation of the *Salmonella* Containing Vacuole (SCV) (Cirillo et al., 1998; Jennings et al., 2017). SPI-2 gene expression is regulated by the two component system SsrAB, which consists of a sensor histidine kinase (SsrA) and its cognate response regulator (SsrB) (Garmendia et al., 2003). Expression and activation of SsrAB is initiated by stimuli in the SCV, most notably acidic pH and low magnesium concentration (Deiwick et al., 1999; Nikolaus et al., 2001; Rappl et al., 2003; Löber et al., 2006). Upon phosphorylation by SsrA, SsrB transcriptionally activates all structural and functional SPI-2 components as well as other promoters outside the SPI-2 locus (Garmendia et al., 2003; Tomljenovic-Berube et al., 2010). SPI-2 is necessary for STm pathogenicity, as its loss prevents intracellular replication and systemic infection (Ochman et al., 1996; Shea et al., 1999). However, the observations of phenotypic heterogeneity in SPI-2 expression by us and others (Blair et al., 2013; Cirillo et al., 1998; Hautefort et al., 2003; Stapels et al., 2018; Reuter et al., 2021; Pospíšilová et al., 2025) suggest a cost to expression that this heterogeneity offsets.

This heterogeneity in SPI-2 expression is distinct from that associated with SPI-1, a separate T3SS and set of secreted effectors STm uses to promote invasion of the gut epithelium. SPI-1 expression is bistable, with a defined fraction of SPI-1(+) and SPI-1(-) cells, and expression imposes a clear growth burden on SPI-1(+) cells relative to their SPI-1(-) counterparts (Ackermann et al., 2008; Saini et al., 2010; Sturm et al., 2011; Diard et al., 2013; Arnoldini et al., 2014; Sánchez-Romero & Casadesús, 2018). In contrast, SPI-2 subpopulations are not defined; individual cells continuously switch into a SPI-2 expressing state over extended periods of time and the rate of this switching is probabilistically tuned to the strength of the intracellular-like stimuli experienced (Spratt & Lane, 2025). These regulatory differences, coupled with the dramatically different environments in which the two virulence systems are required, suggest that SPI-2 phenotypic heterogeneity evolved under distinct selective pressures from SPI-1. However, it remains unknown what costs, if any, make SPI-2 heterogeneity beneficial and why this environment-tunable regulatory strategy evolved.

To address these questions, here we evaluate whether SPI-2 expression incurs a fitness cost in defined *in vitro* environments. We observe that microcolony growth is negatively correlated with single-cell SPI-2 reporter expression. We find increasing levels of the SPI-2 master regulator SsrB imposes increasing context-specific growth and fitness changes, as well as cell morphology effects. These growth costs require SsrB DNA-binding activity, but not its phosphorylation. Lastly, we find that the growth costs incurred by SsrB expression are observed in the absence of the SPI-2 locus, suggesting that the cost of SPI-2 expression is not derived from the T3SS or associated effector proteins themselves, but rather through SsrB-mediated regulation of genes outside of the pathogenicity island. Together, our results reveal an environment-specific cellular burden imposed by SPI-2 expression inherent to the activity of its master regulator SsrB, suggesting that co-regulation of genes involved with cell physiology and growth have shaped the regulatory logic of SPI-2 expression.

## RESULTS

### Single-cell SPI-2 expression is negatively correlated with microcolony growth

Given our prior observations of bimodal expression of SPI-2 (Spratt & Lane, 2025), we hypothesized that expression may be associated with reduced growth at the single-cell level, as has been shown for SPI-1 (Sturm et al., 2011). We attempted to test this in the same way the SPI-1 growth deficit was originally identified, by agarose pad imaging of cells in inducing conditions. We used our previously published plasmid-based reporter for SPI-2 gene expression that consists of a representative SPI-2 promoter (P*ssaG*) fused to sfGFP destabilized with an LVA degradation tag, as well as a constitutive promoter driving mRuby2 expression (Figure S1A, Methods). Surprisingly, we found that we were unable to induce SPI-2 reporter activation using agarose pads made with minimal magnesium, pH 5 media (MgM-MES), despite this media being commonly used to induce SPI-2 (Beuzón et al., 1999; Yu et al., 2004), and having observed strong induction of this reporter in liquid MgM-MES culture in our previous studies. To circumvent this, we grew cells in MgM-MES for four hours to induce SPI-2 in a fraction of the population. We then spotted cells onto agarose pads made with MgM-MES and imaged microcolony formation for 4 hours. Single-cell SPI-2 reporter intensities remained bimodal throughout these experiments (Figure S1B). We observed variation in both SPI-2 reporter expression and colony size (Figure 1A). We therefore classified colonies based on their mean fraction of SPI-2(+) cells across all time as ‘Stable’ (>0.9, ∼30% of colonies), ‘Variable’ (all others, <0.9 and >0.1, ∼25% of colonies), or ‘Negative’ (<0.1, ∼44% of colonies) (Figures 1B and S1C). For each colony, we measured the mean SPI-2 reporter intensity per cell across time, finding that it broadly tracked with final colony cell count (Figure 1B, grey heatmap), consistent with recent findings using a different SPI-2 reporter (Pospíšilová et al., 2025). We also found that expression of SPI-2 was not necessarily stable on agar pads, as evident by colony averages that drop after showing high levels of SPI-2 reporter intensity (Figure 1B). We evaluated whether the SPI-2 expression state at the onset of imaging dictated final colony size, finding a significant size decrease in colonies that started from a SPI-2(+) cell relative to a SPI-2(-) cell, but considerable variation within the SPI-2(-) colonies (Figure 1C). We confirmed that the variation in colony size was not sfGFP(LVA) dependent using a promoterless control, observing bimodal distributions of colony size in the absence of sfGFP(LVA) expression (Figure S1D). Given the fluctuations we observed in SPI-2 reporter expression over the duration of the imaging, we also evaluated whether the mean colony intensity across time related to final colony size, finding a significant negative correlation between the two (Figure 1D). Combined, these results suggest an inverse relationship between single-cell SPI-2 expression and growth rate and motivated further quantification of the fitness costs associated with SPI-2 expression.

**Figure 1.**
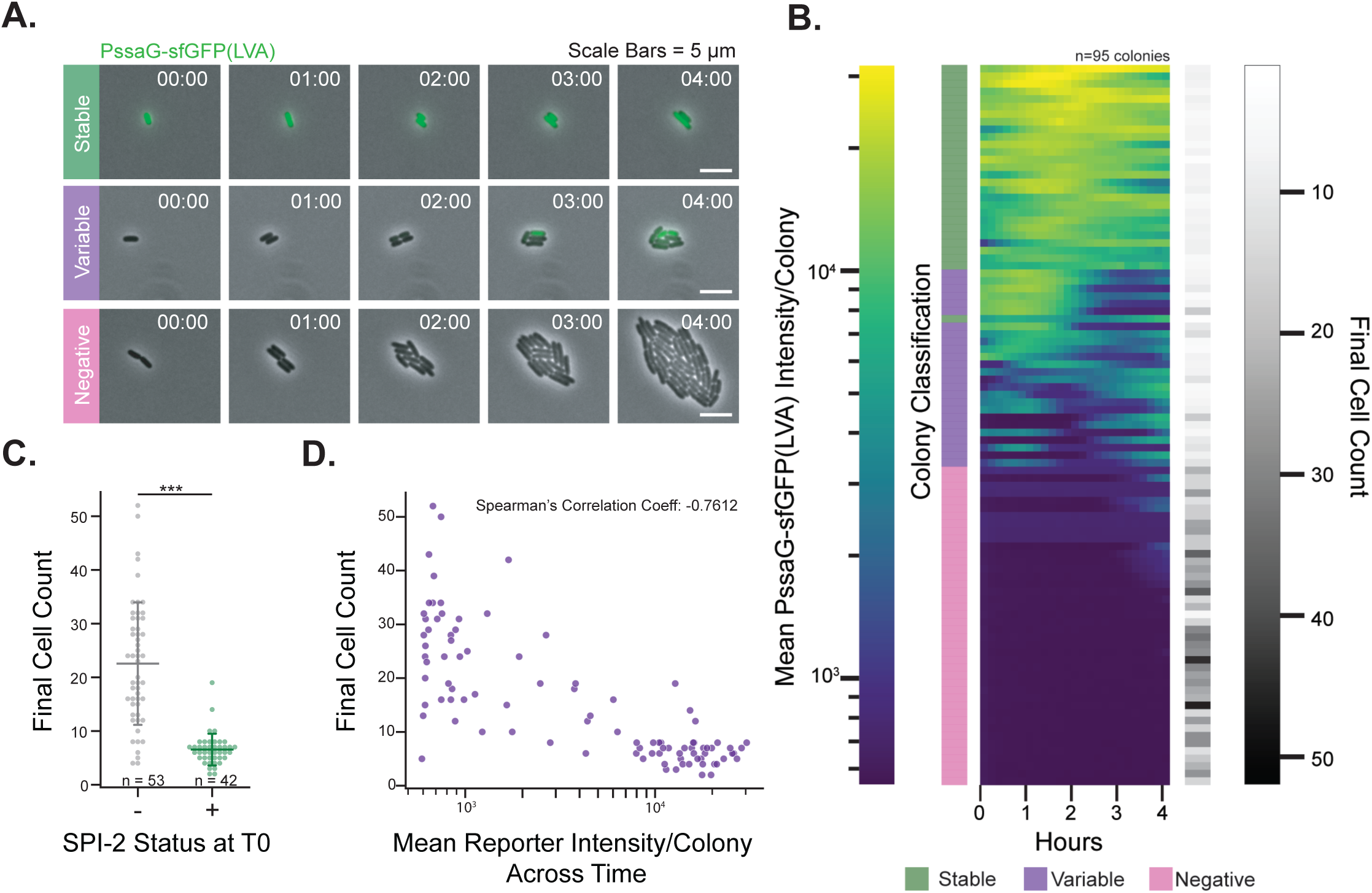
SPI-2 reporter expression is negatively correlated with growth on agarose pads. To evaluate the growth of single cells expressing SPI-2, WT cells containing a SPI-2 reporter (PssaG-sfGFP(LVA)-mRuby2) were grown in SPI-2 inducing (MgM-MES) media for four hours, then spotted onto MgM-MES agarose pads and imaged for four hours. (A) Representative images from one experiment of microcolony growth of cells, with the SPI-2 reporter shown in green, phase in gray. Colonies were classified as ‘Stable’, ‘Variable’, and ‘Negative’ based on their mean fraction of SPI-2(+) cells across time (see Methods and Figure S1C). (B) Heatmap of mean SPI-2 reporter intensity per colony across time, with each row representing a colony and each column a time. The heatmap is sorted by mean colony intensity across all time. Colony classification based on SPI-2 expression is displayed to the left of the heatmap. To connect SPI-2 expression to growth, we measured the final cell count per colony, and a corresponding greyscale single timepoint heatmap showing this value is displayed to the right of the heatmap. (C) Final colony cell count based on SPI-2 reporter status at the beginning of imaging, *** p <0.0001 (Welch’s independent t-test). Dots represent individual colonies. (D) For each colony, the final cell count is plotted against its mean reporter intensity over the time course, showing a negative correlation (Spearman’s correlation coefficient = -0.7612, p<0.001). Data in B-D are pooled from 3 independent experiments, n=95 colonies.

### Expression of the SPI-2 master regulator *ssrB* imposes environment-specific growth and fitness costs

Interpretation of the relationship between SPI-2 expression and growth from these experiments is complicated by both technical and biological factors. We have previously shown that SPI-2 induction is switch-like, committed, and continuously activated at the single-cell level under constant exposure to intracellular-like stimuli in culture and during growth in a mother machine (Spratt & Lane, 2025). The prevalence of ‘Variable’ SPI-2(+) colonies, combined with the inability to robustly induce SPI-2 using agarose pad composition alone, suggests that there are as yet poorly understood environmental factors impacting SPI-2 expression during growth on MgM-MES agarose pads. Further, the need to pre-induce cells prior to imaging makes it impossible to determine whether slowed growth is a cause or a consequence of SPI-2 expression.

To circumvent these issues, we evaluated the fitness costs of SPI-2 expression at the population level by synthetically controlling levels of *ssrB*, the SPI-2 master regulator (Figure 2A - i). This approach also eliminates any concerns with the effects of the SPI-2 reporter on cell physiology and fitness (Figure S1D). We have previously shown that deletion of *ssrB* eliminates downstream SPI-2 gene expression as expected, while constitutive *ssrB* expression from a plasmid (P*_ssrB_*) leads to unimodal SPI-2 reporter expression in both inducing and non-inducing conditions (Spratt & Lane, 2025). To vary SPI-2 expression levels in a controlled, stimuli-independent manner we created a panel of *ssrB* expression plasmids that differ in copy number and the synthetic promoters used (Figure 2A - ii), allowing us to compare strains with no SPI-2 expression (Δ*ssrB*), endogenous regulation of SPI-2 (WT), and three defined levels of *ssrB* ectopic expression (Δ*ssrB* complemented with P_Low-*ssrB*_, P*_ssrB_*, and P_High-*ssrB*_). We generated equivalent constructs expressing sfGFP as plasmid burden controls and to evaluate relative protein production across the panel of plasmids (Figure S2.1A-B).

**Figure 2.**
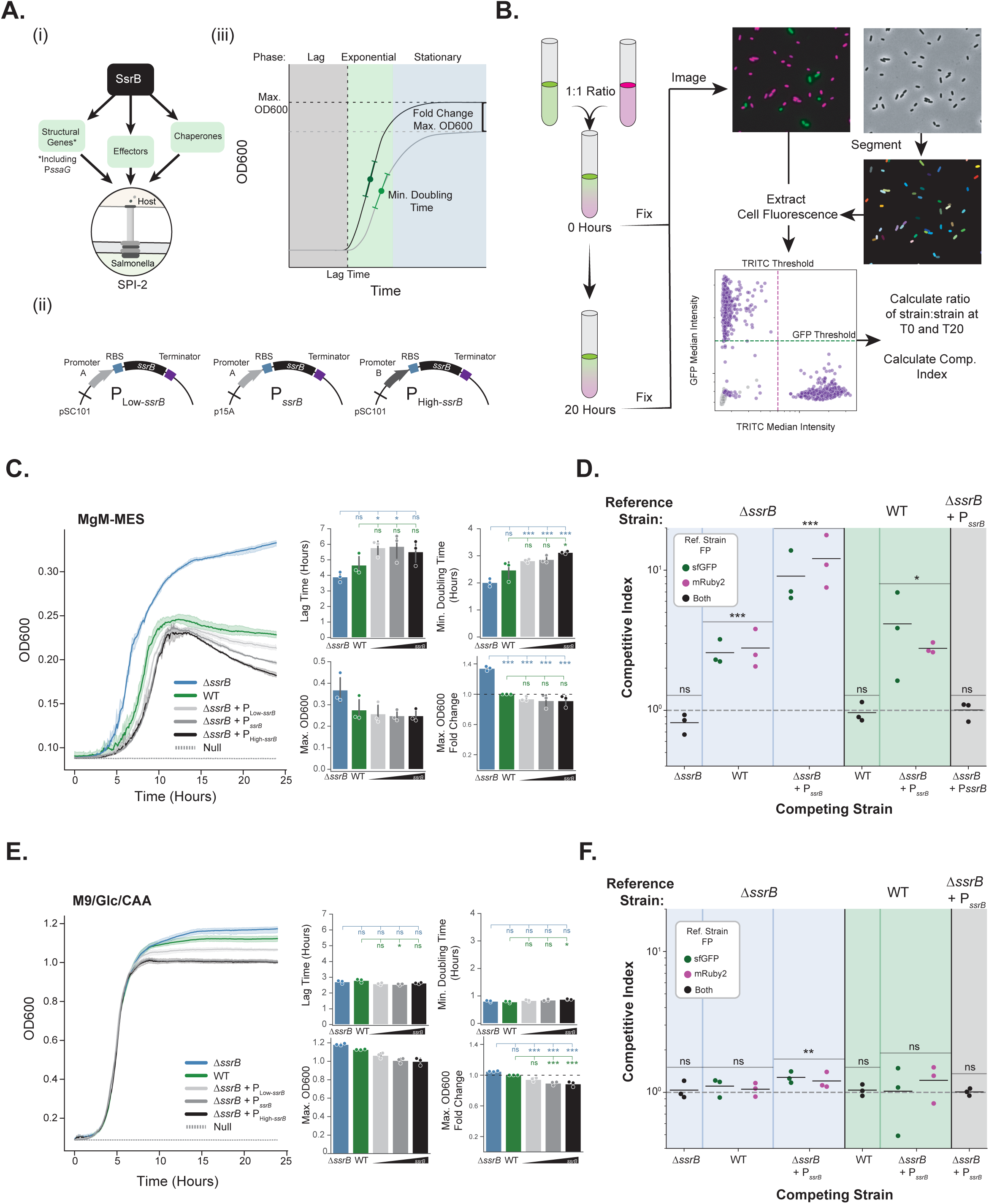
Environment-specific growth and fitness deficits incurred through *ssrB* expression. (A) Varying master regulator *ssrB* expression levels to evaluate fitness costs of SPI-2 expression. (i) The response regulator SsrB is responsible for transcriptionally activating all SPI-2 genes (including *ssaG*, the promoter of which is used in Figure 1). (ii) Design of ectopic *ssrB* expression plasmids using plasmid copy number and synthetic promoter strength to modulate SsrB levels. (iii) Standard bacterial growth curve with summary metrics of interest labeled. Fold Change in max. OD600 is relative to the WT strain (i.e. compare black curve to grey). (B) Schematic representing an imaging-based quantification of a competition assay. Strains expressing either mRuby2 or sfGFP are diluted and mixed at a ratio of 1:1, with a T0 control being fixed in parallel, and grown for 20 hours prior to fixation, imaging, and quantification of relative strain ratios at both time points to calculate the competitive index. (C) Representative growth curves of strains with a range of *ssrB* expression levels and media blank (Null) measured for OD600 in MgM-MES. Shading represents a 95% CI for 3 technical replicates. Barplots show summary metrics for 3 independent experimental replicates including lag time, minimum doubling time, maximum OD600, and fold change in the maximum OD600 relative to the WT strain, * p<0.05, **p<0.01, ***p<0.005, ns = not significant (ANOVA with post-hoc Tukey tests, comparisons only to the WT (green) and Δ*ssrB* strain (blue) are shown, all comparisons in Table S1). (D) Relative fitness of the Δ*ssrB*, WT, and Δ*ssrB* + P*_ssrB_* strains was evaluated via the competition assay described in (B). Competitive index equals the measured ratio of the reference strain (Top, colored background corresponds to growth curve colors) to the competing strain (Bottom) at 20 hours divided by the same measured ratio at the 0 hour mixing timepoint. Each dot represents an independent biological replicate. Fluorescent protein switches (dot color) were performed to control for any effects of fluorescent protein expression on growth. One-sample t-tests were used to evaluate significant differences from an expected ratio of 1. *p<0.05, **p<0.01, ***p<0.005, ns = not significant. (E) Representative growth curves and summary metrics as in (C) in M9/Glc/CAA media. (F) Competitive indices are plotted for pairwise competitions as in (D) in M9/Glc/CAA media.

We first evaluated growth (Figure 2A - iii) and fitness (Figure 2B) of the Δ*ssrB*, WT, and *ssrB* overexpression strains in SPI-2 inducing media, MgM-MES. Growth deficits scaled with *ssrB* expression levels throughout the growth curve, including in lag time, doubling time, and most substantially saturation density (Figure 2C). Notably, these differences were observed between the Δ*ssrB* strain and the WT strain, indicating that endogenous *ssrB* expression levels are sufficient to incur these effects, consistent with our agarose pad data. None of the control sfGFP expression constructs imposed any observable growth effects on the WT strain (Figure S2.1C), indicating that the growth cost was not due to plasmid-based protein production. We also confirmed the specificity of this *ssrB* effect to STm by measuring the growth of *E. coli* strain

MG1655 with the Δ*ssrB* expression constructs, finding these plasmids did not cause a general deficit in the same harsh media conditions (Figure S2.1E). We further tested whether these growth deficits imposed a fitness cost in competition experiments (Figures 2B, S2.2). Strains constitutively expressing *ssrB* were outcompeted by both the Δ*ssrB* and WT strains, and the Δ*ssrB* strain likewise outcompeted the WT strain (Figure 2D). Together, these results demonstrate that in SPI-2 inducing conditions there are growth and fitness deficits associated with expression of the SPI-2 master regulator, *ssrB*.

SPI-2 expression is generally restricted to conditions containing intracellular-like signals, suggesting there is a cost to expression in extracellular conditions. Our synthetic expression constructs allowed us to test this by evaluating the effects of constitutive *ssrB* expression in the absence of intracellular stimuli using neutral, rich media (M9/Glc/CAA). We hypothesized that constitutive *ssrB* expression would elicit a general energetic cost much like that observed with SPI-1. Contrary to our expectation, we found that growth of cells constitutively expressing *ssrB* was nearly indistinguishable from Δ*ssrB* and WT strains, with no deficits in lag or doubling time (Figure 2E). In competition experiments, we also observed no significant fitness benefits of *ssrB* relative to WT, and a marginal benefit relative to the constitutive expression strain (Figure 2F). However, we observed a subtle, but significant reduction in saturation density that scaled with *ssrB* expression level, which held true even when comparing Δ*ssrB* with WT (Figure 2E) and was not observed with increasing sfGFP expression in our control experiments (Figure S2.1D). This likely reflects the known induction of the endogenous SPI-2 system in early stationary phase in rich media (Bustamante et al., 2008). Combined, these data suggest that there are environment-specific growth and fitness deficits associated with expression of the SPI-2 master regulator, *ssrB*, and that these *ssrB*-associated growth deficits are only incurred in stressful or resource-limited conditions.

### *ssrB* expression is associated with increased cell length

The saturation density differences, particularly the increasingly negative slopes observed in stationary phase in MgM-MES, prompted us to image cells after 24 hours of growth to evaluate cell morphology. While cell morphology appeared grossly normal across conditions and strains, quantification of cell length revealed a subtle increase with increasing *ssrB* expression levels (Figures 3A-B). This dose-dependent effect was most pronounced in MgM-MES media, though the highest overexpression construct did promote a significant cell length difference in M9/Glc/CAA relative to the WT and Δ*ssrB* strains. The cell length effects were not due to protein expression levels, since the equivalent GFP expression constructs did not affect cell length (Figure S3A). These results indicate that SsrB affects cell growth and physiology beyond its role in promoting expression of SPI-2.

**Figure 3.**
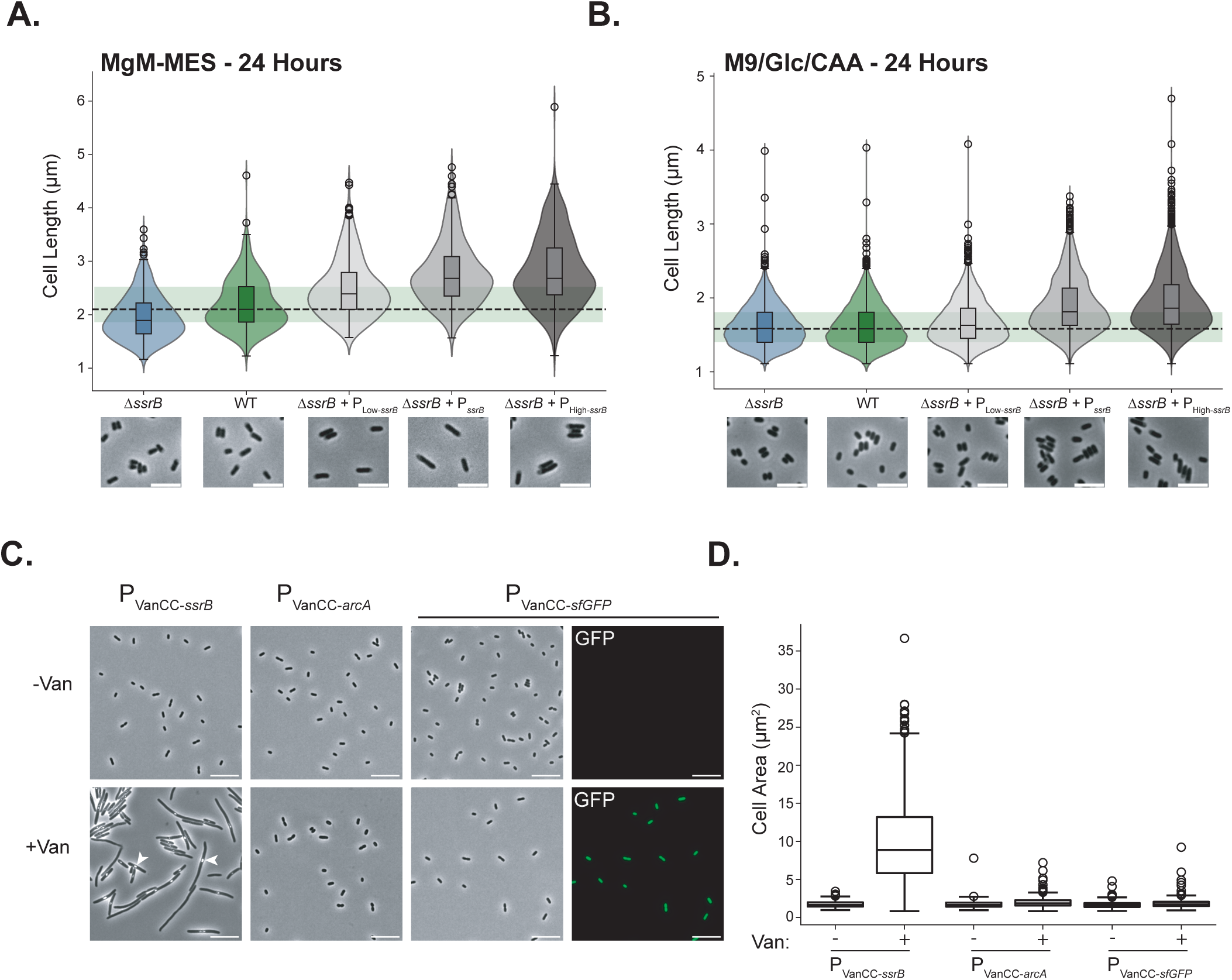
Cell morphology effects observed with *ssrB* expression. (A, B) Strains were imaged following 24 hours of plate reader growth in (A) MgM-MES and (B) M9/Glc/CAA. Cell lengths were measured for individual cells and are displayed as violin plots with overlaid boxplots. Representative images are shown for each strain and condition, scale bars = 10µm. n=450 cells per strain, 150 from each of three independent experiments for MgM-MES, n=4500 cells per strain, 1500 from each of three independent experiments for M9/Glc/CAA. Dashed black line shows the median cell length of the WT strain, light green shading represents the interquartile range of the WT. (C) Representative images of P_VanCC-*ssrB*_, P_VanCC-*arcA*_, and P_VanCC-sfGFP_ after 4 hours of growth with or without inducer (vanillic acid - Van) are shown, scale bars = 10µm. Examples of protein aggregates observed after inducing *ssrB* are highlighted with white arrowheads. GFP channel is shown for P_VanCC-sfGFP_, and GFP intensity images are scaled equivalently. (D) Quantification of cell area for samples shown in (C), n = 1350 cells per condition, 450 sampled from each of 3 independent experiments.

Our attempts to clone higher level constitutive *ssrB* expression constructs were unsuccessful, which we attributed to a potential toxicity effect. Given the subtle cell morphology effects we observed at the tolerated expression levels, we hypothesized that this toxicity might be linked to *ssrB-*mediated effects on cell growth and length. To evaluate *ssrB* effects above these low constitutive levels, and to determine whether we could generate a more pronounced cell length phenotype, we constructed an inducible *ssrB* expression plasmid using a vanillic acid inducible promoter/repressor system from the *E. coli* Marionette library (Figure S3B) (Meyer et al., 2019). This system was selected for its dynamic range, tightly repressing gene expression in the absence of inducer while promoting expression far above our constitutive constructs upon induction. As controls, we generated equivalent constructs with sfGFP and *arcA*, another DNA-binding response regulator not associated with SPI-2 expression, to test if morphology effects were *ssrB* specific. Following 3 hours of induction with vanillic acid, we fixed and imaged strains (Figure 3C). We observed that *ssrB*-expressing cells displayed a severe filamentation phenotype, with cell areas on average 10-fold larger than control strains expressing sfGFP or *arcA* (Figure 3D). Notably, these morphology effects were specific to *ssrB*. Vanillic acid induces sfGFP expression to high levels (high enough that imaging cannot be used to quantify its expression relative to the constitutive plasmids used in Figure S2.1), with minimal leaky expression in the uninduced conditions as expected (Figure 3C and S3C). Inducible sfGFP expression did not cause any noticeable changes in cell growth (Figure S3D) or cell size. While overexpression of *arcA*, an endogenous response regulator with myriad promoter targets like *ssrB*, affected cell growth as expected (Figure S3D), it did not produce a filamentation or substantial cell morphology phenotype. Taken together with the subtle cell length increases observed in stationary phase, these results directly implicate *ssrB* in STm cell morphology regulation, an unexpected finding given its primary role in regulating SPI-2 expression, and further demonstrate the sensitivity of STm cell physiology to increasing SsrB levels.

### SsrB-mediated growth deficits require DNA binding, but not phosphorylation

We next investigated whether SsrB-driven growth deficits were dependent on its DNA-binding activity or phosphorylation status. SsrB consists of a receiver domain (amino acids 1-149) and a DNA-binding domain (residues 152-212) (Figure 4A (Abramson et al., 2024; Goddard et al., 2018)). DNA binding is facilitated by phosphorylation of aspartate 56 by the sensor kinase SsrA, which induces a conformational change that exposes alpha helix 3 of the DNA-binding domain, allowing for dimerization and subsequent binding of target gene promoters. While previous work has shown that overexpression of SsrB alone is sufficient to activate downstream SPI-2 genes, their expression still requires a phosphorylation at D56, suggesting phosphorylation may occur through other donors (Feng et al., 2004). To evaluate the contribution of phosphorylation, DNA-binding, and dimerization we generated plasmids expressing previously characterized SsrB mutants associated with each of these functions, D56A, K179A, and L201A, respectively (Figure 4A). We complemented the WT and Δ*ssrB* strains with these constructs and compared their growth in MgM-MES (Figure 4B). We expected all three mutations to fully rescue the growth deficit incurred by P*_ssrB_* and phenocopy the Δ*ssrB* strain. This was true of both mutations in the DNA-binding domain (K179A, L201A), indicating growth deficits are the direct result of SsrB binding DNA and not a consequence of general toxicity due to overexpression. In contrast, the phosphosite D56A mutant only partially rescued growth, ultimately failing to recapitulate Δ*ssrB* level growth. As a complementary approach to investigate the role of D56 phosphorylation, we evaluated the growth of strains lacking the SsrA kinase in combination with ectopic SsrB expression (Figure S4A). The Δ*ssrAB* strain grew virtually identically to the Δ*ssrB*, and when complemented with the P*_ssrB_* had a partial rescue of only saturation, similar to that of the D56A mutants.

**Figure 4.**
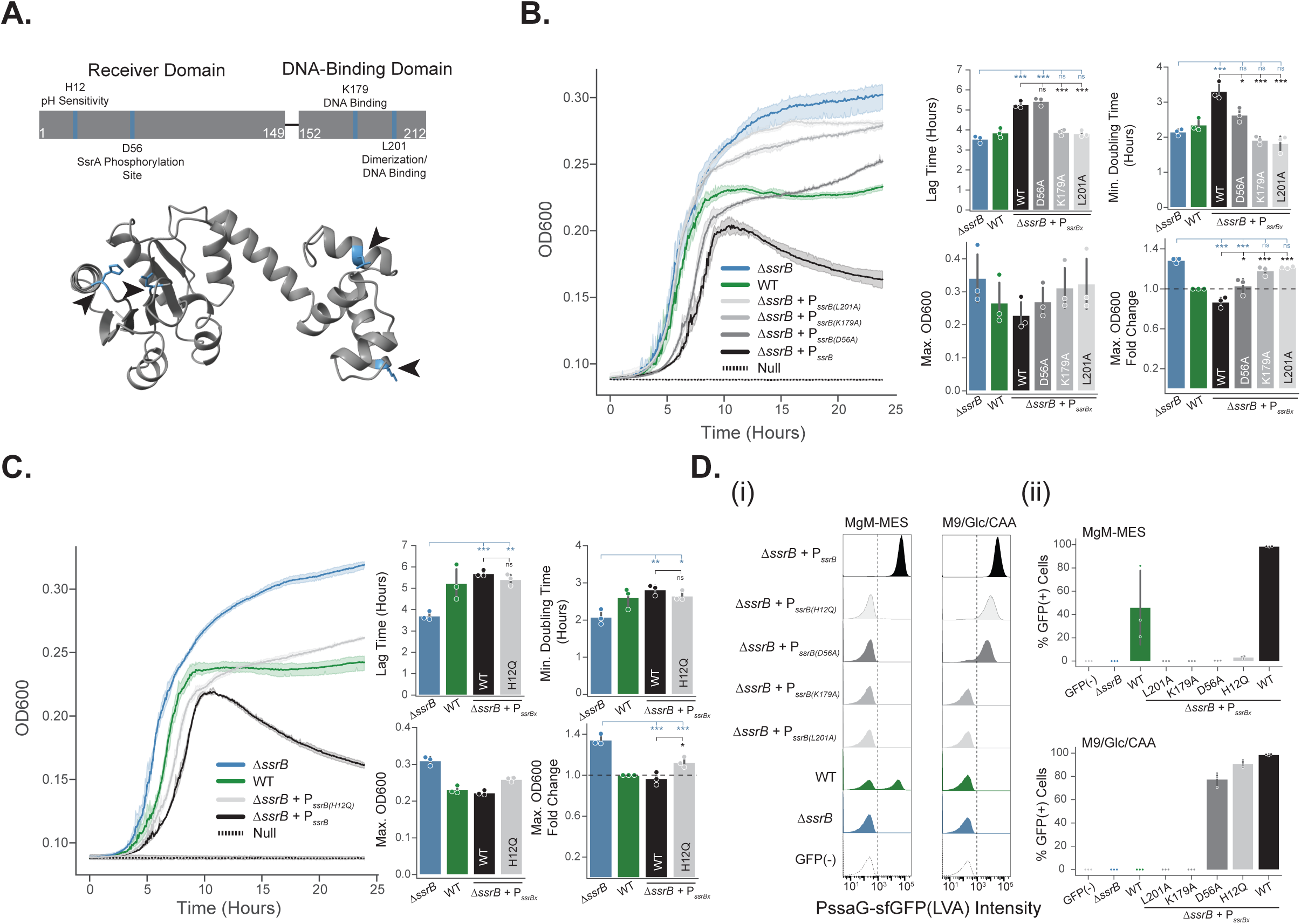
Growth effects of constitutive expression of *ssrB* mutants. (A) Top - Schematic representation of the 2 major domains of the SsrB protein, with key residues (blue lines) and their functional significance depicted. Bottom - AlphaFold rendered structure of WT SsrB with residues of interest displayed as stick models in blue, black arrowheads for emphasis. (B) Representative growth curves showing the effects of the K179A, L201A, and D56A SsrB mutations in MgM-MES. OD600 was measured for 24 hours, error shading represents 95% CI of technical triplicates, media null control is shown as dashed line. Summary metrics from 3 independent experiments are displayed, error bars = standard deviation. * p<0.05, **p<0.01, ***p<0.005, ns = not significant (ANOVA with post-hoc Tukey tests, comparisons only to the Δ*ssrB* strain (blue) and Δ*ssrB*+P*_ssrB_* (black) are shown, all comparisons in Table S1). (C) Representative growth curves showing growth effects of the H12Q SsrB mutation in MgM-MES. OD600 was measured for 24 hours, error shading represents 95% CI of technical triplicates. Summary metrics from 3 independent experiments are shown, error bars = standard deviation. p-values are annotated as in (B), full comparisons in Table S1. (D) SPI-2 reporter expression in strains expressing WT or mutant SsrB. All strains contain the SPI-2 reporter, except for the GFP(-) strain, which contains a promoterless control plasmid. Strains were grown in MgM-MES or M9/Glc/CAA for four hours, fixed, and analyzed via flow cytometry. Representative histograms of SPI-2 reporter intensity are shown, dashed line = GFP(+) threshold, n = 50,000 cells per sample (i). Summary of SPI-2 reporter (+) cells for 3 independent experiments are shown (ii).

Beyond its receiver domain phosphorylation and DNA-binding functions, SsrB has also been shown to undergo pH-dependent conformational changes that increase DNA binding efficiency. Given that we observe *ssrB*-mediated growth costs in pH 5 MgM-MES and in M9/Glc/CAA stationary phase, in which mild acidification occurs, we hypothesized that these growth costs may be dependent on this conformational switch. We therefore evaluated the role of histidine residue 12, which has been linked to this pH-dependent enhancement of SsrB DNA binding upon acidification of the bacterial cytoplasm. The mutation H12Q has been shown to disrupt this conformational change, and reduce phosphorylation (Liew et al., 2019). Overexpression of this mutant led to a partially rescued growth phenotype, similar to that observed in the phosphosite D56A mutant, indicating that this acid-dependent conformational change is not required to cause the growth deficits (Figure 4C).

We were surprised that mutations affecting phosphorylation and pH sensitivity failed to relieve the growth cost of SsrB expression, since these features are critical for SPI-2 gene expression at endogenous SsrB levels (Walthers et al., 2007). We therefore evaluated how ectopic constitutive expression of these mutants impacted SPI-2 reporter output relative to the WT SsrB protein. To do so, we co-transformed the Δ*ssrB* strain with each SsrB expression construct and the PssaG-sfGFP(LVA)-mRuby2 reporter (Figure S1A). We then used flow cytometry to measure reporter output in both SPI-2 inducing (MgM-MES) and non-inducing conditions (M9/Glc/CAA) (Figure 4D). Consistent with our previous results, overexpression of WT SsrB resulted in unimodal, high-intensity SPI-2 reporter activity in both conditions, while the WT strain showed no signal in M9/Glc/CAA and bimodal reporter intensity after 4 hours in MgM-MES, and the Δ*ssrB* strain showed no signal in either condition. Both DNA-binding domain mutations (K179A and L201A) eliminated SPI-2 reporter output in both conditions. The phosphorylation and pH sensitivity (D56A and H12Q) mutants retained unimodal, although lower intensity, SPI-2 reporter activity in M9/Glc/CAA, consistent with SPI-2 activation due to the expected accumulation of SsrB in early stationary phase. In MgM-MES however, these mutants showed almost no SPI-2 reporter activity after 4 hours of growth, likely because in this condition SPI-2 reporter output depends more on signaling-dependent post-translational modification of SsrB, while protein accumulation is expected to be negligible at this timepoint. Our results indicate that the DNA-binding mutants eliminate SPI-2 gene expression while the phosphorylation mutants cause a substantial reduction. However, this reduction was insufficient to fully rescue growth in MgM-MES. Further, while P*_ssrB_* expression in the Δ*ssrAB* strain showed higher SPI-2 reporter expression than the D56A and H12Q mutants (Figure S4B), likely a result of SsrA-independent phosphorylation of SsrB, it did not show a corresponding growth deficit (Figure S4A). Taken together, these data suggest that the growth cost of SsrB expression is disconnected from SPI-2 gene expression.

### SsrB-mediated growth deficits are SPI-2 independent

Our original hypothesis was that SsrB-dependent growth deficits were due to the energetic costs associated with SPI-2 gene expression and T3SS production. However, the cell morphology effects associated with increasing *ssrB* expression and the incongruency of growth deficits relative to SPI-2 reporter output across the SsrB mutant proteins suggested that SsrB may impair fitness through SPI-2 independent mechanisms. To directly test whether the SsrB-driven activation of SPI-2 is responsible for growth costs, we deleted the entire 25kb SPI-2 locus, spanning *ssrA* to *ssaU*. All of the structural components of the T3SS and the majority of effectors are encoded here, although several effector operons are located outside this locus. As expected, the ΔSPI-2 strain grew nearly identically to the Δ*ssrB* strain in MgM-MES (Figure 5A). However, when complemented with P*_ssrB_*, the strain did not phenocopy the ΔSPI-2 strain.

**Figure 5.**
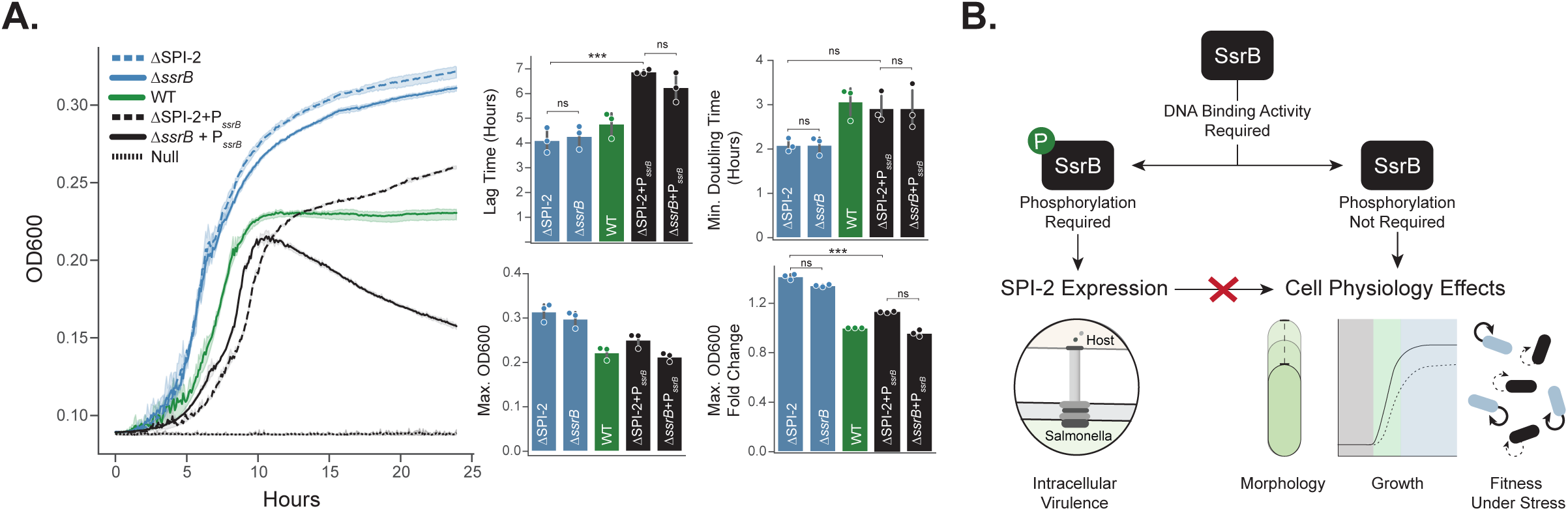
*ssrB* mediated growth deficits are T3SS independent. (A) Representative growth curves evaluating the effects of a SPI-2 locus deletion, error shading represents 95% CI of technical triplicates. Summary metrics from 3 independent experiments are shown, error bars = standard deviation, * p<0.05, **p<0.01, ***p<0.005, ns = not significant (ANOVA with post-hoc Tukey tests, all comparisons in Table S1). (B) Summary of findings. The environment-specific effects of *ssrB* expression on cell physiology that we report are DNA-binding-dependent, but phosphorylation-independent. Cell physiology effects are seen in the absence of the SPI-2 locus, meaning that SsrB regulated expression of the T3SS and secreted effectors occurs in parallel with, but independently from, the SsrB-dependent cell physiology phenotypes.

Instead, there was a partial growth rescue relative to P*_ssrB_* in the Δ*ssrB* strain, similar to both the phosphosite mutant and the Δ*ssrAB* strain (Figures 4B, S4A). This indicates that the effect of the full SPI-2 deletion on growth was due to the loss of the SsrA kinase, and hence the reduced phosphorylation of SsrB. From this, we conclude that the costs of SPI-2 expression are not due to energetic or cellular resource burdens of T3SS assembly or effector secretion. Instead, our results demonstrate that the growth and fitness costs are driven by SsrB regulated targets outside of the SPI-2 locus.

## DISCUSSION

While phenotypic heterogeneity in virulence factor expression has been broadly observed, direct evaluations of the fitness costs that this heterogeneity is often assumed to offset remain limited. The initial aim of this work was to characterize and quantify the fitness costs of SPI-2 expression following our observation that single-cell SPI-2 reporter activity negatively correlates with growth. To this end, we used constitutive ectopic expression of the SPI-2 response regulator *ssrB* to deterministically control SPI-2 expression, revealing environment-specific deficits in growth, fitness, and cell morphology that correlated with SsrB levels. We ultimately uncovered, however, that the growth burden of SPI-2 expression is not due to the production of the T3SS itself, but rather the DNA-binding activity of its master regulator on targets outside the SPI-2 locus (Figure 5B). These results have implications for understanding the evolution of SPI-2 regulation and the impact of SsrB activity during infection, but also highlight the necessity of directly quantifying virulence burdens following observations of phenotypic heterogeneity.

The T3SS-independent nature of the fitness deficits distinguishes SPI-2 from other virulence systems in which the end product itself imposes a metabolic or energetic burden (Septer et al., 2023; Greene et al., 2025). The SsrB target genes that are responsible for these fitness and morphology effects remain unknown. ChIP-seq and microarray data that elucidated the SsrB binding motif identified numerous binding sites outside of the SPI-2 locus (Tomljenovic-Berube et al., 2010). However, none of these are obvious candidates to explain the growth deficits incurred by ectopic SsrB expression. RNA-seq experiments in the specific conditions we have associated with differential fitness (MgM-MES, late stationary phase) and with our constitutive expression constructs may better elucidate candidate target genes.

Our observations that phosphorylation of SsrB is not required to elicit SsrB-mediated growth deficits suggest that transcription of these unknown target genes may involve unphosphorylated SsrB. Phosphorylation has been shown to affect SsrB target promoter affinity, with unphosphorylated SsrB preferentially targeting biofilm associated genes and phosphorylation mediating a lifestyle transition to virulence gene expression (Desai et al., 2016). Alternatively, the target genes responsible for the growth deficits may simply have a higher binding affinity for or sensitivity to unphosphorylated SsrB than SPI-2 genes. This would be consistent with the fact that in our ectopic system SsrB is produced at high enough concentrations that activating signals are not needed for SPI-2 reporter expression, and our observation of marginal rescues when phosphorylation capacity is reduced or eliminated.

While the mechanism behind SsrB-associated growth costs remains unclear, the environment-specific nature of the costs provides insight into the potential benefits of heterogeneous expression of SPI-2. Although the SPI-2 T3SS does not directly impose growth deficits in the *in vitro* conditions tested, its production requires SsrB, meaning that in the endogenous system virulence and cell physiology effects are inherently coupled. Our previous work has shown that increasing strength and integration of intracellular signals accelerates the rate at which individual cells transition into a SPI-2 expressing state. SsrB being costly in acidic, nutrient-limited non-phagosomal environments, therefore, supports a model in which the population benefits from probabilistic tuning of SPI-2 in ambiguous inducing environments.

Alternatively, the SsrB targets outside of the SPI-2 locus may confer a fitness advantage in the intracellular environment while imposing a deficit extracellularly. While it is unlikely this fitness benefit arises from slowed growth alone, it may be advantageous for cells to replicate more slowly in the phagosomal environment, limiting host cell death and subsequent inflammation. Recent work has further proposed that bifurcation of intracellular STm into SPI-2 expressing and non-expressing subpopulations reflects cooperative virulence, in which SPI-2(+) cells secrete and grow slowly, while SPI-2(-) cells benefit and promote outgrowth (Pospíšilová et al., 2025). Heterogeneous expression of SPI-2 may therefore confer multiple context specific benefits: limiting the cost of expression in harsh or ambiguous extracellular environments, while enabling division-of-labor during intracellular infection.

Our work further uncovers divergent selective pressures acting on the two T3SSs in STm, SPI-1 and SPI-2, likely reflecting their distinct environments and functions. For example, the environment-dependent nature of the SPI-2 growth cost is in contrast to SPI-1, which has been shown to cause growth deficits even in rich media when overexpressed (Cooper et al., 2017; Sturm et al., 2011). One explanation for this difference may be that SPI-1 imposes a higher resource burden on cells; injectisomes per cell have been measured at 50-100 for SPI-1, and only 1-5 for SPI-2 (Chakravortty et al., 2005; Kubori et al., 2000). One notable commonality between the two virulence factors is that the SPI-1 growth cost is also only partially rescued through the deletion of the secretion apparatus (Sturm et al., 2011). This suggests that SPI-1 regulators may drive growth costs through parallel pathways similar to what we have observed for SsrB. Such regulator-derived fitness effects may prove more widespread across virulence systems; disentangling the fitness consequences of virulence regulators from their end products is therefore critical to understanding how selective pressures have shaped regulatory mechanisms.

Combined, these results clarify the selective pressures acting on SPI-2 by identifying environment-specific growth and fitness costs that are driven by the expression and DNA-binding activity of the response regulator SsrB rather than the T3SS itself. These findings highlight the need to directly quantify and assess specific virulence factor costs to gain insight into the functional benefits of the evolved regulatory mechanisms.

## MATERIALS AND METHODS

### Strains, Plasmids, and Growth Conditions

*Salmonella enterica* serovar Typhimurium (ATCC strain 14028s) was obtained from the lab of Michael McClelland and was used as the WT strain in all experiments. MG1655 *E. coli* was obtained from the *E. coli* Genetic stock center. The Δ*ssrB* strain was used from the STm knockout library (Porwollik et al., 2014). Δ*ssrB* was PCR confirmed using a primer located outside the coding region of the deleted gene and a primer for the chloramphenicol resistance cassette replacing the gene, or a primer within the open reading frame of the deleted gene.

Where indicated, bacteria were cultured in M9 minimal medium (1X M9 Salts with 0.1mM CaCl_2_, 2mM MgSO_4_, 0.4% glucose) supplemented with 0.2% casamino acids (M9/Glc/CAA) or else pH 5.0 MgM-MES (170mM 2-[N-morpholino] ethane-sulphonic acid (MES) adjusted to pH 5.0 with NaOH, 5mM KCl, 7.5 mM (NH_4_)_2_SO4, 0.5mM K_2_SO_4_, 1mM KH_2_PO_4_, 8µM MgCl_2_) supplemented with 38mM glycerol and 0.1% casamino acids (Beuzón et al., 1999; Yu et al., 2004).

Multi-gene knockouts (*ssrAB* and SPI-2) were generated using a lambda red recombination based approach identical to the one described in (Porwollik et al., 2014). Briefly, we transformed WT STm with plasmid pKD46 that contains the lambda red proteins under control of an arabinose-inducible promoter. This plasmid further contains a temperature-sensitive origin of replication, such that growth at 37°C will elicit loss of the plasmid. For all steps preceding recombination, these cells were grown at 30°C.

A chloramphenicol resistance cassette was PCR amplified with primers that added 40bp of homology to target sequences within the *ssrA* and *ssrB* genes (for Δ*ssrAB*) or within *ssrA* and *ssaU* (for ΔSPI-2). Target sequences were located 40bp within the ORFs of interest to minimize effects of the insertion. Following transformation of PCR products into WT STm containing pKD46, cells were grown shaking for 4 hours at 37°C to allow recombination to occur and then plated onto LB agar with chloramphenicol to select for the successfully integrated cassette. Selected colonies were whole genome sequenced to confirm the deletion and sequencing results analyzed with Breseq software (Deatherage & Barrick, 2014).

Plasmids were constructed using the EcoFlex library for golden gate cloning of transcriptional units (Moore et al., 2016). The custom promoter/repressor fragment used for the VanCC inducible constructs was PCR amplified from the *E. coli M*arionette library (Meyer et al., 2019). The *ssrB* open reading frame was PCR amplified from WT STm. All *ssrB* point mutation constructs were made by PCR-based incorporation of the nucleotide base pair changes from the WT P*_ssrB_* template followed by Gibson assembly into the EcoFlex Level 1 plasmid. Plasmid components and sources are listed in Table S2. For strains containing plasmids, all experiments were performed with selective antibiotic for plasmid maintenance unless otherwise noted.

To prepare electrocompetent STm strains, a glycerol/mannitol step centrifugation protocol as described in (Warren, 2011) was used. Briefly, a culture of the given strain was grown to exponential phase, then pelleted at 2000 g at 4°C for 15 minutes and resuspended in ice-cold water. A solution of 1.5% mannitol and 20% glycerol was then slowly dispensed into the cell suspension. Cells were then pelleted with gradual acceleration and deceleration at 2000g at 4°C for 15 minutes. Supernatant was aspirated and cells were resuspended in glycerol/mannitol solution between 250 and 500x their concentration in the original culture. Following competent cell preparation, 100 ng of plasmid was transformed into the strain via electroporation at 1800 kV, 200 Ω, 25 µF. To generate strains containing two plasmids, competent cells were prepared of the strain containing one plasmid and then transformed with the second plasmid.

To make glycerol stocks of strains 700µL of an overnight culture grown in LB with required antibiotics was added to 300µL of 50% w/v glycerol, vortexed lightly and stored at -80°C.

### Plate Reader Growth Curves and Analysis

Cultures were inoculated from a single colony and grown in M9/Glc/CAA for 16-18 hours. From these stationary phase cultures, 1mL of culture was pelleted and resuspended in PBS. This resuspension was then diluted to an OD of 0.02 in the media to be used in the plate reader.

Cultures were grown for 4 hours, then diluted to an OD of 0.0015265 in fresh media of interest. This dilution was then pipetted in 150µL aliquots into 3 wells of a clear bottom 96-well plate.

Well plates were sealed with adhesive plastic film, and holes were made in the top of each well with a 30 gauge hypodermic needle to allow for aeration. Cells were grown in a Tecan Spark Microplate reader. Following a settling time of 15 minutes to equilibrate temperature, OD600 was recorded every 5 minutes for 24 hours, with orbital rotation speed of 240 rpm. Growth curve data was analyzed and plotted using the Omniplate package (Montaño-Gutierrez et al., 2022).

### Fluorescence Microscopy

Images were acquired using a Nikon Ti2-E microscope with Lumencor Spectra III light engine, Prime BSI CMOS camera, motorized stage, perfect focus system, and Oko environmental enclosure with temperature control and 2x2 binning. Imaging data used in plots is available in Table S3, all single cell data is unfiltered and not downsampled.

### Agar Pad Live-Cell SPI-2 Induction

Cultures of WT STm containing the PssaG-sfGFP(LVA) or the promoterless reporter control (GFP(-)) were inoculated from single colonies and grown for 16-18 hours in 3mL of M9/Glc/CAA. Cells were then diluted 1:100 into fresh M9/Glc/CAA and grown for 4 hours. OD600 was measured and cells were diluted to an OD of 0.02 in MgM-MES media. Cells were grown for 4 hours to induce SPI-2.

During growth, a 2% w/v agarose pad was created by melting agarose in MgM-MES media. Liquid agarose/MgM-MES was pipetted onto an ethanol cleaned coverslip. A second coverslip was set on top of the first, creating an agarose ‘sandwich’. The agarose sandwich was allowed to dry for 45 minutes before being cut into approximately 1cm^2^ pieces.

After four hours of growth in MgM-MES, cells were diluted to an OD of 0.05 in MgM-MES. 2 µL of this dilution was spotted onto the agarose pieces and allowed to dry for 3-5 minutes. Agarose pads were then flipped onto a warm LabTek single chamber slide. A wet kimwipe was added to the corner of the chamber to help retain moisture in the pad. The chamber was wrapped in parafilm, and then imaged.

Images were captured every 10 minutes for a total of 8 hours. Images were acquired using 30ms exposure for TRITC and 50 ms exposure for GFP. All replicates were captured at 50% LED power for sfGFP and TRITC, with the exception of 1 replicate in which 30% LED power was used for TRITC alone; as TRITC intensity was only used for segmentation which was unaffected by the lower power, this replicate was included. Images were manually cropped to contain single colonies. Colonies were selected for further analysis based on focal consistency and lack of cell overlap within the first 4 hours. Cropped images were segmented from the mRuby2 signal using Omnipose (Cutler et al., 2022) with the pretrained ‘bact_fluor_omni’ model, a flow threshold of 0, and a mask threshold of 2. Segmentation was manually corrected as needed using Napari. Intensity measures of individual cell labels were extracted using sci-kit regionprops. SPI-2(+) threshold was set as 3 times the mean intensity of the GFP(-) colonies from all experiments. This threshold was applied consistently across replicates.

### Post-Plate Reader Imaging and Analysis

Imaging for GFP intensity (Figure S2) and cell length (Figures 3A and B) was performed immediately following plate reader experiments displayed in Figure 2 and S2. Aliquots of plate reader culture were spotted onto 2% agarose pads made with PBS and imaged within 30 minutes of experiment conclusion. Phase images were captured at 500ms and GFP images were captured at 70ms at 50% LED power.

Images were segmented from the Phase channel using Omnipose with the pretrained ‘bact_phase_omni’ model, a flow threshold of 0, and a mask threshold of 1.5. Cell labels were filtered on the basis of area (between 40 and 250 pixels, minimum size adjusted down due to small size after 24 hours growth), eccentricity (>=0.6), and solidity (>=0.8) to exclude debris. Cell length (as ‘max_feret_diameter’) and sfGFP median intensity were extracted using sci-kit regionprops.

### Culture Competition Assays and Analysis

Designated strains were transformed with both the P_Red_ and P_Green_ expression constructs independently (plasmids were assembled via EcoFlex Golden Gate cloning as described above, components in Table S2). Cells were prepared equivalently to the plate reader preliminary dilutions. After four hours of growth following the first dilution, cells were diluted to a final OD of 0.0015625 in 3mL of media (either M9/Glc/CAA or MgM-MES) and combined in a 1:1 ratio for all pairwise comparisons. At the same time, 600µL of total culture was fixed in the same volumetric ratio as the tubes were prepared in order to normalize to the initial mixture of cells (T0). Cells were fixed by adding 200µL of 16% PFA to the 600µL of culture, inverting to mix, incubating at room temperature for 30 minutes, and then washing twice with PBS. Samples were stored in a final mL of PBS at 4°C until imaging. For all pairwise comparisons, a fluorescent protein flip was also performed. All strains were competed with their own opposing fluorescent protein counterpart as a control. Cells were grown in pairwise mixtures for 20 hours, and then fixed as for T0. All samples were imaged within a week of fixation. Fixed cells were imaged on 2% agarose/PBS pads with 500ms Phase exposure, 50ms GFP exposure, and 50ms of TRITC exposure, with both LEDs at 50% power.

Images were segmented in the Phase channel using Omnipose with the ‘bact_phase_omni’ model, a flow threshold of 0, and a mask threshold of 1.5. Labels were filtered on the basis of area (between 50 and 250 pixels), eccentricity (>=0.6), and solidity (>=0.8) prior to any fluorescence-based filtering. Each timepoint and media had fluorescent thresholds set independently. Thresholds were set conservatively to exclude fluorescent protein negative cells and cells falling outside of the bulk of the fluorescent protein positive populations. Thresholds were set the same across experiments. Rare instances of “dual positive” events were removed from further analysis.

Cells defined as GFP or TRITC positive were then used to calculate ratios of Reference Strain: Competing Strain for the 0 hour and 20 hour timepoints. The competitive index for each strain was calculated as the 20 hour ratio divided by the 0 hour ratio.

### Induced Filamentation Experiments and Analysis

A 1 mM vanillic acid stock solution was prepared in ethanol. Cultures of STm strains were inoculated from single colonies and grown in M9/Glc/CAA for 16-18 hours. Cultures were back diluted 1:100 into 3mL of fresh M9/Glc/CAA and grown for 3 hours. OD600 was measured and cultures were again diluted into 3mL of M9/Glc/CAA either with or without 1µM vanillic acid.

Cultures were grown in inducing conditions for four hours, then fixed as per the protocol described in the competition assay. All samples were imaged within 1 week of fixation. Images for the filamentation assay VanCC-sfGFP control were captured at 10ms exposure and 50% LED power for GFP. All images were captured at 500ms for Phase.

All images were segmented in the phase channel using the Omnipose ‘bact_phase_omni’ model. For non-filamentous samples, the same Omnipose settings were used as in the competition assays. For filamentous samples, the parameters were adjusted to a mask threshold of -4.2 and a flow threshold of 0. Cell shape parameters and sfGFP median intensity were extracted using sci-kit regionprops. Filters for cell shape and size were adjusted to be less conservative, such that filamentous cells would not be excluded from the dataset, with only objects of area less than 50 pixels, eccentricity less than 0.5, and solidity less than 0.6 removed. Cell length (‘max_feret_diameter’) was not measured due to the curvature of the filamentous cells.

### SPI-2 Reporter *in vitro* Induction and Flow Cytometry

Strains containing the PssaG-sfGFP(LVA)-mRuby2 were started from isolated colonies and grown overnight in M9/Glc/CAA for 16-18 hours. Overnight cultures were back-diluted to an OD of 0.02 in fresh M9/Glc/CAA and grown for four hours. After four hours growth, cells were back diluted to an OD of 0.02 in MgM-MES, and a 600µL sample of culture was also fixed for the M9/Glc/CAA sample. Cells were grown for four hours in MgM-MES, then fixed as per the protocol described in the competition assay. Flow cytometry was performed within one week of fixation.

Fixed cells were diluted into PBS such that they could be analyzed at approximately 10,000 events per second or below. Samples were analyzed using a BD LSRFortessa SORP Cell Analyzer with HTS (6-laser 18-parameter). At least 50,000 mRuby2 positive events per sample were captured.

Analysis and plotting of flow histograms was done using FlowJo version 10. Gates were drawn for each experiment and are consistent across samples within an experiment. Because of the presence of the mRuby2 signal in all reporter samples, light scatter gates were drawn liberally to encompass the bulk of the densest region of events. To eliminate debris events, mRuby2 positive samples were then identified by drawing a gate such that less than 0.5% of the negative control sample was classified as mRuby2 positive. mRuby2 positive events were then down-sampled to exactly 50,000 events using the FlowJo plugin DownSample (ver. 3.3.1). All further analyses and plotting were performed on down-sampled, mRuby2 positive events. The %GFP(+) cells (Table S4), were exported from FlowJo and plotted using custom Python code, Seaborn (version 0.13.1) and Matplotlib (version 3.8) packages.

## Supporting information

Reagents_Table

## ACKNOWLEDGEMENTS

We thank the members of the Lane Lab for their sharing of protocols, code, and valuable feedback on this work. We thank the Robert H. Lurie Comprehensive Cancer Center of Northwestern University for the use of the Flow Cytometry Core Facility, which provided training and equipment for flow cytometry experiments. The Lurie Cancer Center is supported in part by an NCI Cancer Center Support Grant #P30 CA060553. We acknowledge our funding sources: NIH Cellular and Molecular Basis of Disease Training Program award to MRS (NIH T32 GM008061).

---

**Figure S1:**
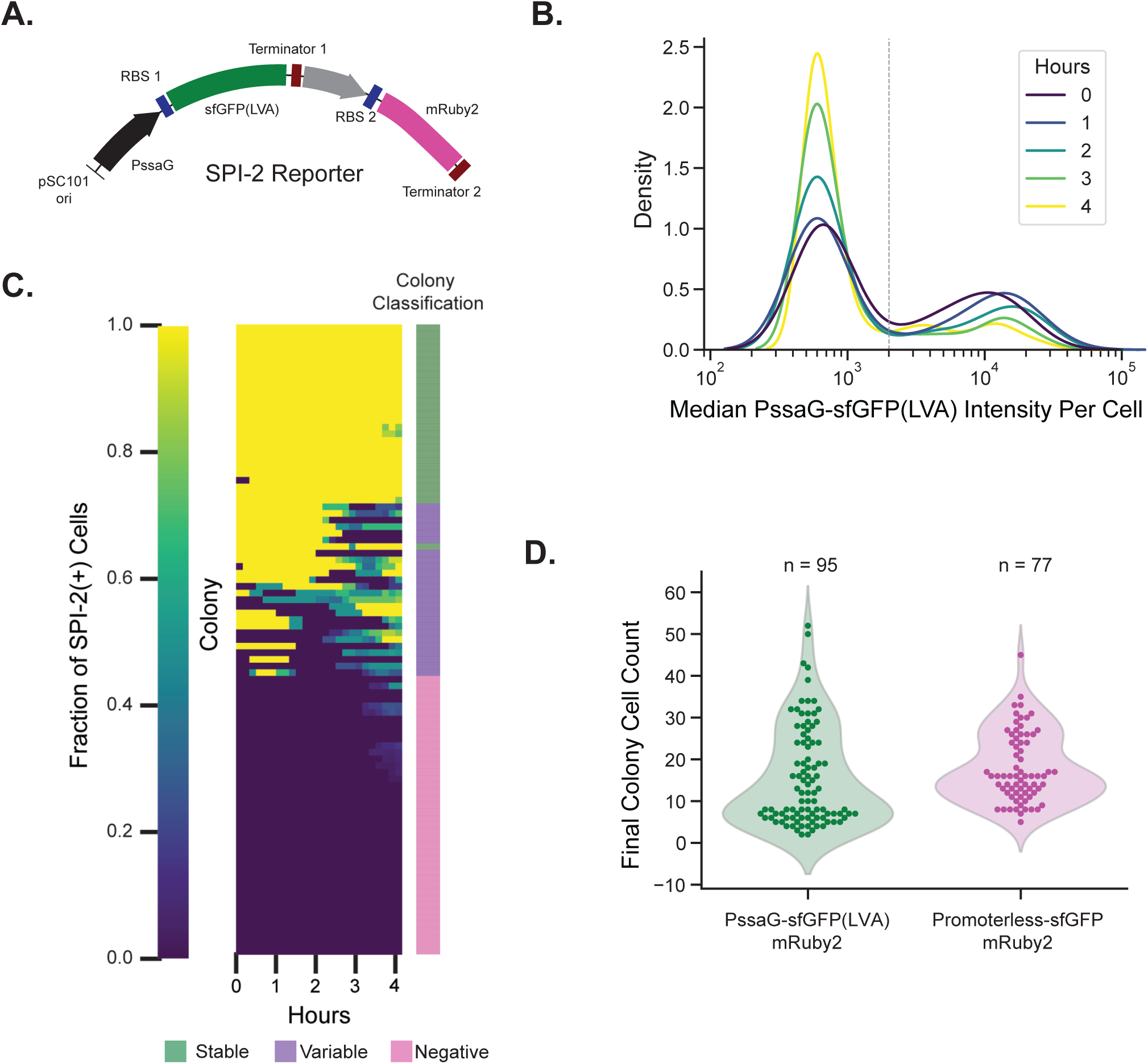
(A) Schematic of the major components of the SPI-2 reporter. (B) Distributions of single-cell SPI-2 reporter intensities after 0, 1, 2, 3, and 4 hours growth on agarose pads, respectively. Data is pooled from 3 independent experiments, n = 132, 246, 429, 743, and 1470 cells. The grey dashed line represents the threshold used to classify cells as SPI-2(+) (3 times the mean of the promoterless-sfGFP control pooled from all three experiments). (C) Heatmap of fraction of cells per colony classified as SPI-2(+) based on the threshold in (B). Cells with a mean SPI-2(+) fraction across time greater than 0.9 are classified as ‘Stable’ (green), less than 0.1 are classified as ‘Negative’ (pink), and all others are classified as ‘Variable’ (purple). (D) Final colony cell count distributions for strains containing the SPI-2 reporter or the promoterless sfGFP control. Although the distributions differ, both show bimodality in final colony cell count.

**Figure S2.1:**
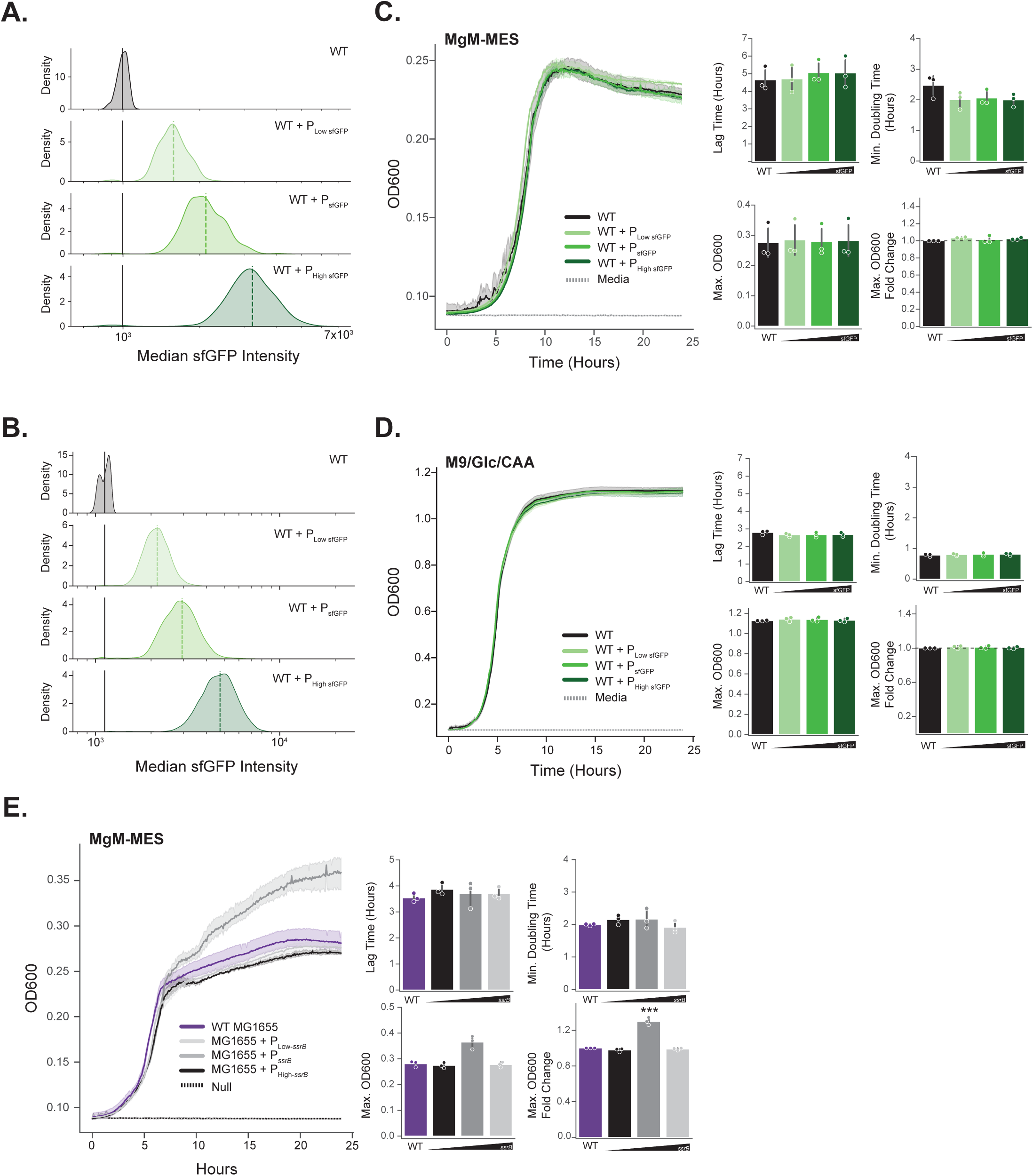
sfGFP overexpression plasmids equivalent to the P*_ssrB_* plasmids in Figure 2 were used to show that growth deficits observed due to *ssrB* expression are not because of plasmid burden and that relative protein production levels are distinct. (A-B) Density plots of single-cell sfGFP intensity as measured by imaging at the conclusion of the plate reader experiments in MgM-MES (A) or M9/Glc/CAA (B). KDE plots for MgM-MES n = 450 cells (150 from each of 3 experiments), KDE plots for M9/Glc/CAA n = 4500 cells (1500 from each of 3 experiments). Mean of WT (GFP-) cells is shown as a solid black line. Means for the respective plasmids are shown as dashed green lines. (C-D) sfGFP overexpression does not cause any observable growth differences relative to WT STm. Representative growth curves of WT STm containing the sfGFP plasmids in MgM-MES (C) or M9/Glc/CAA (D), error shading represents 95% CI of technical triplicates. Summary metrics are shown for 3 independent experiments for each media condition, no statistically significant differences (ANOVA with Tukey’s, all comparisons in Table S1). (E) Growth costs of *ssrB* expression are *Salmonella* specific. Representative growth curves of *E. coli* MG1655 with three levels of *ssrB* overexpression constructs grown in MgM-MES. Shading represents a 95% CI for 3 technical replicates. Summary metrics from 3 independent experiments are shown, ***p<0.005 (ANOVA with Tukey’s compared to all groups, full comparisons in Table S1)

**Figure S2.2:**
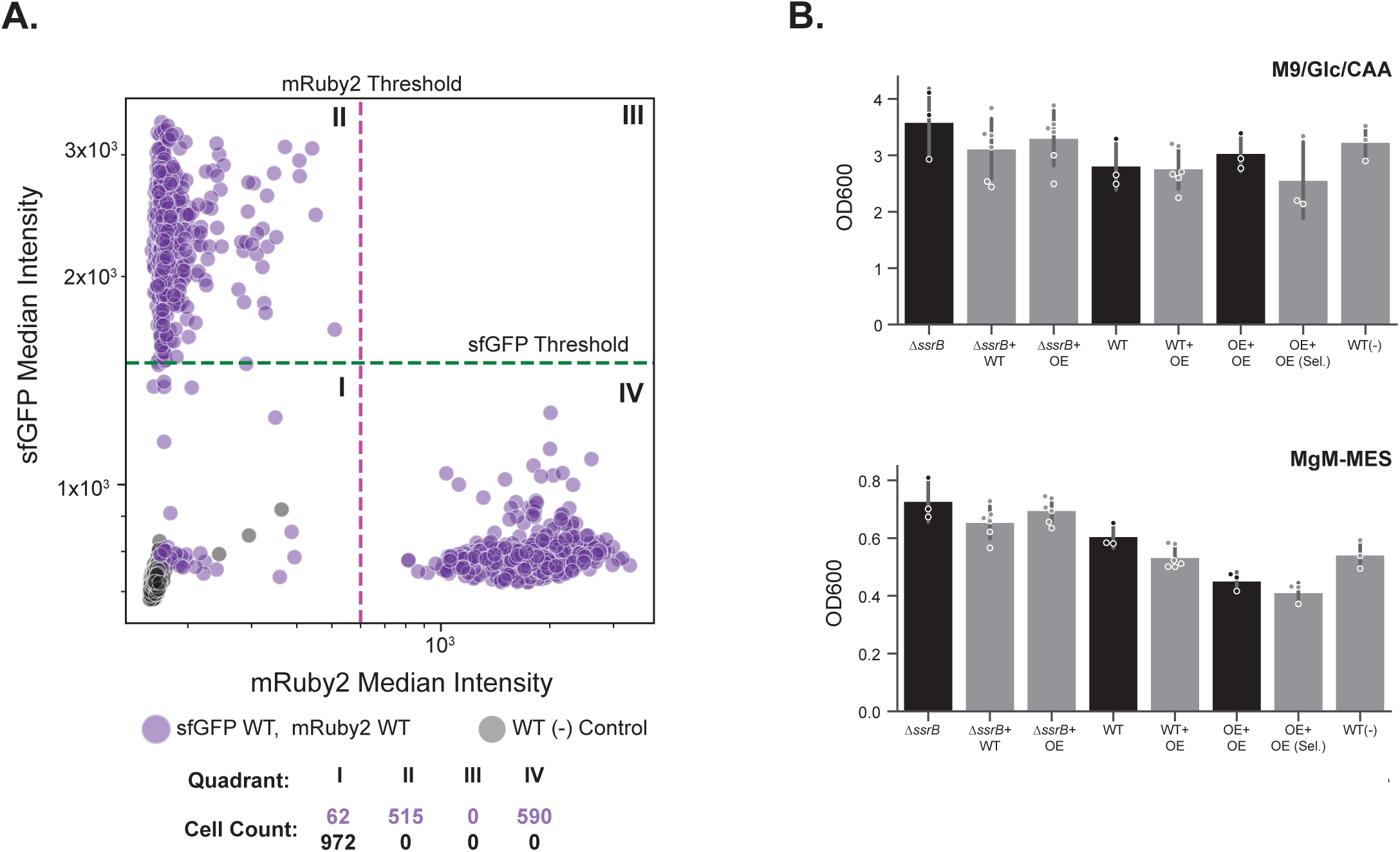
(A) Example thresholding of samples for the competition assay. Thresholds were set conservatively to exclude fluorescent protein negative cells and cells falling outside of the bulk of the fluorescent protein positive population. Thresholds were set independently for media and time conditions, but were applied consistently across experiments. Samples shown are the WT plasmid-free control (WT(-) Control, gray) and a mixed sample of WT expressing sfGFP or mRuby2 at T0 (purple). Cell counts within each quadrant for the two strains are displayed. (B) OD600 readings were measured at the 20 hour timepoint following competition assays. OE = overexpression: Δ*ssrB*+P*_ssrB_*, OE+sel indicates strains grown with P*_ssrB_* plasmid-selecting antibiotics for comparison. While variable, saturation OD600 trends with *ssrB* expression as expected, with mixed populations growing to densities generally between those of the two strains grown independently.

**Figure S3.**
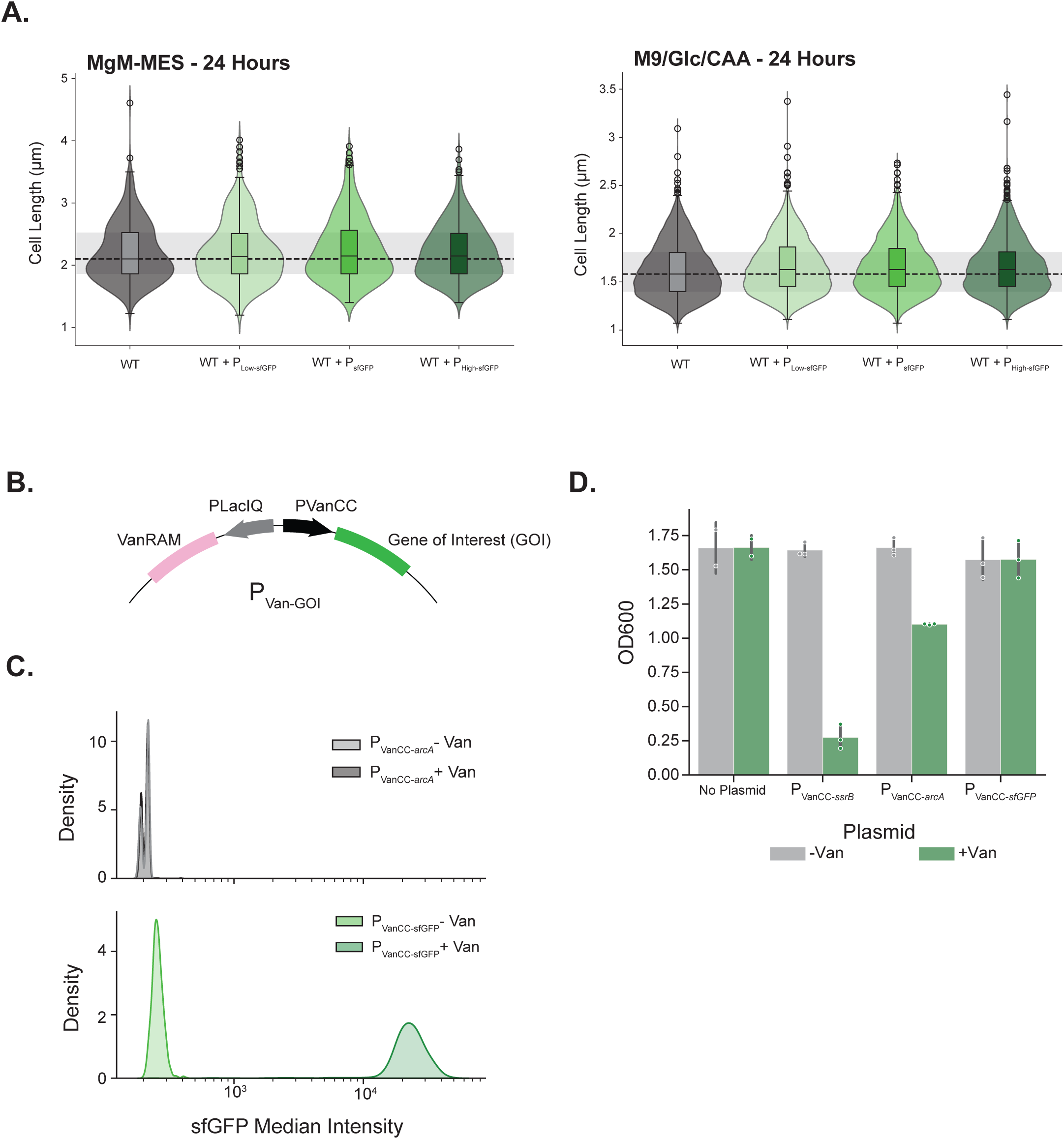
(A) Plasmid-expression of sfGFP does not affect cell length. Cell length of strains containing the sfGFP expression constructs are shown after growth in the plate reader for 24 hours in the listed conditions. (B) Schematic of the VanCC inducible system including the repressor (VanRAM) under control of the constitutive PLacIQ and the gene of interest under control of the VanCC promoter. Upon addition of vanillic acid, the repressor is released from the VanCC promoter and transcription is activated. (C) Median single-cell sfGFP intensity of STm containing P_VanCC-sfGFP_ with and without inducer as measured from images in Figure 3C, n = 1350 per condition, 450 sampled from each of 3 independent experiments. P_VanCC-*arcA*_ with and without inducer is displayed as a fluorescence negative control. GFP(-) peaks (P_VanCC-*arcA*_ -Van) are variable day to day due to minor background differences, producing distinct peaks when replicates are combined. (D) OD600 readings were measured from cultures after growth in media with or without vanillic acid inducer for 4 hours. Induction imposes a growth deficit in P_VanCC-*arcA*_ and P_VanCC-*ssrB*_.

**Figure S4.**
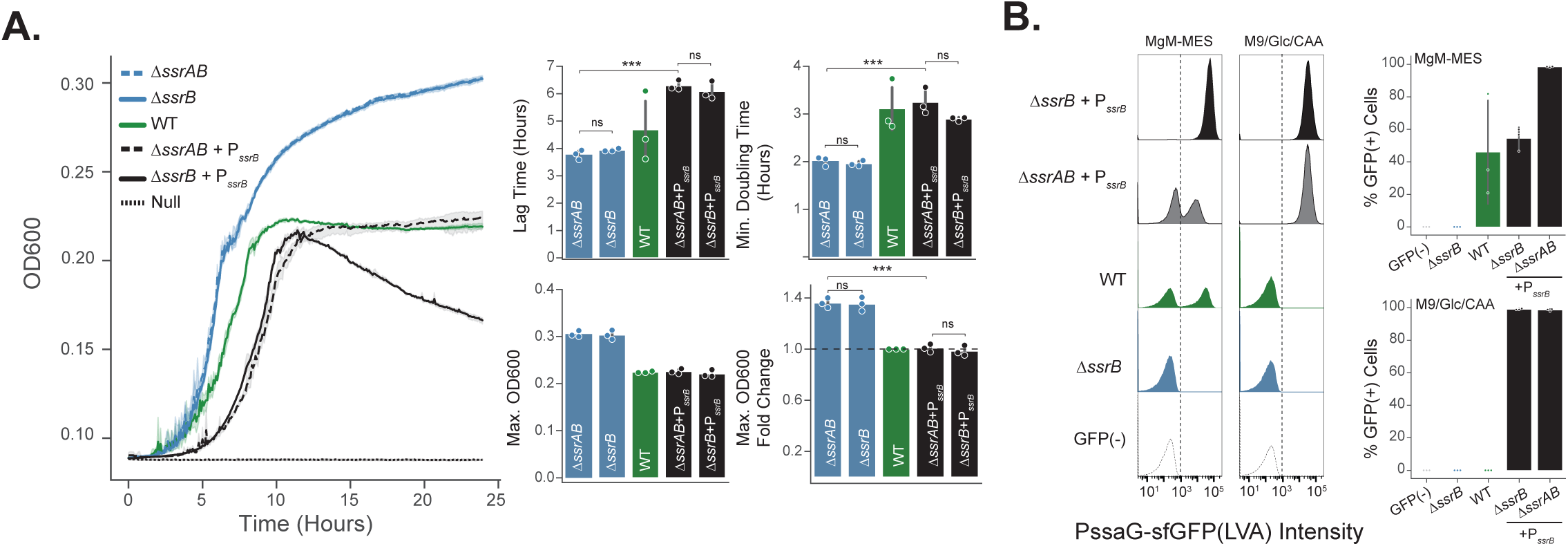
(A) Strains constitutively expressing *ssrB* do not phenocopy the Δ*ssrB* strain in the absence of the sensor kinase SsrA. Representative growth curves of Δ*ssrAB* strains with and without P*_ssrB_* compared to controls in MgM-MES, error shading represents 95% CI of technical triplicates. Summary metrics for equivalent growth curves from 3 independent experiments are shown, error bars = standard deviation, * p<0.05, **p<0.01, ***p<0.005, ns = not significant (ANOVA with post-hoc Tukey tests, all comparisons in Table S1). (B) SPI-2 reporter output in the Δ*ssrAB* and Δ*ssrB* strains complemented with P*_ssrB_*. Histograms are from one representative experiment, n = 50,000 cells per sample. All control samples are from the same experiments as in Figures 4D.

## Notes

### Competing Interest Statement

The authors have declared no competing interest.

