## Supplementary material for "Regulator-derived growth and fitness costs of *Salmonella* SPI-2 expression are environment specific and T3SS independent": Reagents_Table

| REAGENT or RESOURCE | SOURCE | IDENTIFIER |
| --- | --- | --- |
| **Bacterial and virus strains** | | |
| *Salmonella enterica* serovar *typhimurium* (STm) | McClelland Lab | ATCC 14028 |
| STm Strain 14028 Δ*ssrB* (Chlor^R^) | Porwollik S. et al. (2014) | N/A |
| STm Strain 14028 Δ*ssrAB* (Chlor^R^) | This study | N/A |
| STm Strain 14028 ΔSPI-2 (Chlor^R^) | This study | N/A |
| **Chemicals, peptides, and recombinant proteins** | | |
| M9 Salts | Fisher Scientific | Cat# DF0485-17 |
| Acid Casein Peptone (CAA) | Fisher Scientific | Cat# BP1424 |
| MES Hydrate | Sigma-Aldrich | Cat# M2933 |
| Electron Microscopy Sciences 16% Paraformaldehyde Aqueous Solution, EM Grade | Fisher Scientific | Cat# 50-980-487 |
| Research Products International Corp, Low Melt Temperature Agarose | Fisher Scientific | Cat# 50-213-122 |
| **Recombinant DNA** | | |
| PssaG-sfGFP(LVA)-mRuby2 | Spratt & Lane (2025) | N/A |
| Ppromoterless-sfGFP-mRuby2 | Spratt & Lane (2025) | N/A |
| P_Low-_*_ssrB_* | This paper | N/A |
| P*_ssrB_* | Spratt & Lane (2025) | N/A |
| P_High-_*_ssrB_* | This paper | N/A |
| P_Low-sfGFP_ | This paper | N/A |
| P_sfGFP_ | This paper |  |
| P_High-sfGFP_ | This paper |  |
| P*_ssrB(D56A)_* | This paper | N/A |
| P*_ssrB(H12Q)_* | This paper |  |
| P*_ssrB(K177A)_* | This paper |  |
| P*_ssrB(L201A)_* | This paper |  |
| See Table S2 for all plasmid components |  |  |
| EcoFlex MoClo kit | Addgene | Kit # 1000000080 |
| pKD46 | ECGRC | CGSC 7739 |
| **Software and algorithms** | | |
| FIJI | Schindelin, J. et al. (2012) | https://fiji.sc/ |
| Omnipose | Cutler K. et al. (2022) | https://github.com/kevinjohncutler/omnipose |
| Omniplate | Montaño-Gutierrez et al. (2022) | https://git.ecdf.ed.ac.uk/pswain/omniplate |
| Python scripts for data analysis and plotting | This paper | N/A |
| AlphaFold3 | Abramson, J et al. (2024) | https://alphafoldserver.com |
| ChimeraX | Goddard TD, et al. (2018) | https://www.cgl.ucsf.edu/chimerax/ |
| **Other** | | |
| Microplate with Lid, 96 Well, Black | Tecan | Cat# 30122306 |
| Microplate, 96 Well, μClear®, Black | Greiner | Cat# 655097 |
